# Dynamic modelling of the PI3K/mTOR signalling network uncovers biphasic dependence of mTORC1 activation on the mTORC2 subunit Sin1

**DOI:** 10.1101/2020.11.30.403774

**Authors:** Milad Ghomlaghi, Sungyoung Shin, Guang Yang, David E. James, Lan K. Nguyen

## Abstract

The PI3K/mTOR signalling network critically regulates a broad array of important biological processes, including cell growth, metabolism and autophagy. Dysregulation of PI3K/mTOR signalling is associated with a variety of human diseases, including cancer and metabolic disorders. The mechanistic target of rapamycin (mTOR) is a kinase that functions as a core catalytic subunit in two physically and functionally distinct complexes termed mTOR complex 1 (mTORC1) and mTORC2, which also share other common components such as mLTS8 (also known as GβL) and DEPTOR. Despite being the subject of intensive research, a full picture of how mTORC1/2 assembly and activity are coordinated, and how they are functionally connected remain to be fully characterised. This is due primarily to the complex network wiring, featuring a growing number of intricate feedback loops and post-translational modifications, which require quantitative systems-level approaches to decipher. Here, we integrate predictive computational modelling, *in vitro* experiments and -omics data analysis to elucidate the dynamic and emergent features of the PI3K/mTOR network behavior. We construct new mechanistic models of the network that encapsulate novel critical mechanistic details, including mTORC1/2 coordination by mLTS8 (de)ubiquitination, and Akt-to-mTORC2 positive feedback loop. Model simulations subsequently confirmed by experimental validation revealed a previously unknown biphasic, threshold-gated dependence of mTORC1 activity on the key mTORC2 subunit Sin1, which is robust against cell-to-cell variation in protein expression. Furthermore, our results support the essential role of mLST8 in both mTORC1 and 2 activity, and suggest mLST8 could serve as a viable therapeutic target in breast cancer. Overall, our integrated analyses provide fresh systems-level insights into the dynamic behavior of PI3K/mTOR signalling and shed new light on the complexity of this important network.

**AUTHOR SUMMARY:** Signalling networks are the key information-processing machineries that underpin the ability of living cells to respond proportionately to extra- (and intra-) cellular cues. The PI3K/mTOR signalling network is one of the most important signalling networks in human cells that regulates cellular response to hormones such as insulin, yet our understanding of the network behaviour remains far from complete. Here, we employed a highly integrative approach that combines predictive mathematical modelling, biological experimentation, and data analysis to gain novel systems-level insights into PI3K/mTOR signalling. We constructed new mathematical models of this complex network incorporating important regulatory mechanisms. In contrary to commonly held views that mTORC2 lies upstream and is a positive regulator of mTORC1, we found that their relationship is highly nonlinear and dose dependent. This finding has major implications for mTORC2-directed anti-cancer strategies as depending on the cellular contexts, blocking mTORC2 may reduce or even enhance mTORC1 activation, the latter could inadvertently blunt the effect of mTORC2 blockade. Furthermore, our results demonstrate that mLST8 is required for the assembly and activity of both mTOR complexes, and suggest mLST8 is a viable therapeutic target in breast cancer, notably breast cancer.

## INTRODUCTION

The PI3K/mTOR signalling network plays an important role in the regulation of cell signal transduction and regulates a variety of key biological processes such as cell growth, metabolism and autophagy (1). The mechanistic target of rapamycin (mTOR) is a Ser/Thr kinase that lies at the center of this complex network, where it serves as an indispensable catalytic subunit for two functionally distinct complexes termed mTOR complex 1 (mTORC1) and mTOR complex 2 (mTORC2). In addition to mTOR, mTORC1 and 2 share two common subunits, mLST8 (also known as GβL) and DEPTOR, whereas Raptor and PRAS40 are unique components of mTORC1 (2), and Sin1 (3) and Rictor (4) are exclusive to mTORC2. Reflecting its importance in physiological regulation, the PI3K/mTOR network is frequently disrupted in human diseases, including cancer, metabolic and neurodegenerative diseases (2). In cancer alone, more than 40 inhibitors directed at various components of the network have been developed or are under active development (5). Given the clinical relevance of PI3K/mTOR signalling, it is important to understand the interconnectivities within this network and emergent network behaviors.

The PI3K/mTOR network is highly complicated and arguably one of the most extensively studied signalling pathways, yet its complexity continues to expand through new mechanistic discoveries. For example, in addition to known feedback mechanisms such as S6K-mediated negative feedback to PI3K/Akt via IRS, we have identified a positive feedback loop between Akt and mTORC2, where Akt phosphorylates Sin1 to enhance mTORC2 activity (5, 6). More recently, a molecular switch involving mLST8 through its (de)ubiquitination modification was identified (7). Mechanistically, the TRAF2 E3 ubiquitin ligase promotes mLST8 ubiquitination on Lysine 63 (K63), which disrupts its interaction with the unique mTORC2 component Sin1 (7). By contrast, ubiquitinated mLST8 can be converted back to its de-ubiquitinated form by the OTUD7B deubiquitinase. De-ubiquitinated mLST8 binds more favourably to Sin1, which facilitates mTORC2 assembly but at the same time reduces mTORC1 formation (7) (Fig 1A). These findings add extra layers of complexity and intricacy to the wiring of the PI3K/mTOR network, yet its dynamic properties incorporating these new regulatory mechanisms have not been characterized.

**Fig 1.**
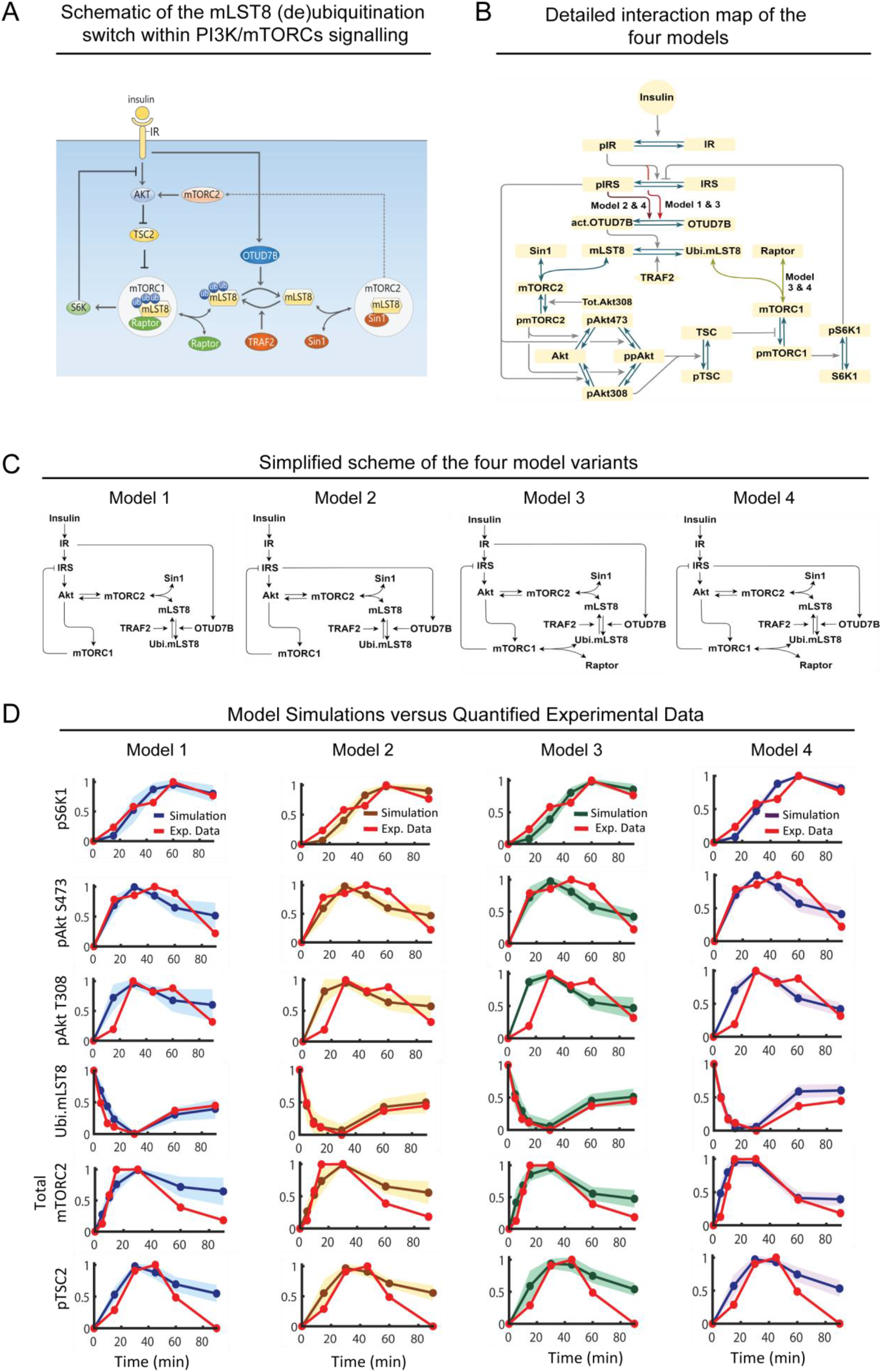
Construction and calibration of new mathematical models of the PI3K/mTOR signalling network incorporating the mLST8 (de)ubiquitination switch. **(A)** Schematic displaying a molecular switch mediated by mLST8 ubiquitination recently reported in (7), which dictates how mLST8 associates with mTORC1 or 2 through Raptor and Sin1, respectively (ubi: ubiquitination). **(B)** Detailed reaction diagrams of the new PI3K/mTOR network models incorporating the mLST8 switch. Four model variants (models 1-4) with distinguishing interactions as described in the text are highlighted (ubi: ubiquitination, act: activated, Tot: total). **(C)** Simplified schematics of the different network structure captured by the four model variants. **(D)** Time-course model simulations using best-fitted parameter sets as compared to the quantified experimental data (red curves), presented for each of the four models. Experimental data represent the dynamic response of various network proteins following insulin stimulation (100nM) in MEF cells, obtained from (7). Per model, multiple best-fitted sets obtained from model calibration (n= 38, 26, 11, 17 for model 1, 2, 3, 4, respectively) were used for simulations: the lines represent the mean simulated curve and the shaded areas indicate standard deviation of all curves.

Moreover, although the mLST8 ubiquitination-dependent switch seems to provide tight regulation of mTOR complex integrity, the functional role of mLST8 in each complex remains to be fully characterised. While it is established that mLST8 is indispensable for mTORC2 activity (8, 9), whether it is essential for mTORC1 function remains unclear. For example, ablation of mLST8 does not affect mTORC1 activity as measured by the phosphorylation level of its major substrate S6K (9-11). On the other hand, mLST8 stabilizes the Raptor-mTOR interaction and promotes mTORC1 activity (12), and upregulation of mLST8 enhances mTORC1/2 activities in human colon carcinoma and prostate cancer (13).

Here we employed an integrated approach that combines predictive network modelling and biological experiments to analyse the emergent network-level behavior of PI3K/mTOR signalling conferred by the mLST8-induced switch. For this, we constructed new mechanistic models of the PI3K/mTOR network that explicitly encapsulate mLST8 (de)ubiquitination, ensuing mTORC1/2 coordination, and the novel Akt-mTORC2 positive feedback. The alternative models consider different competing hypotheses of network interactions, reflecting different network structures. To distinguish among the possible model variants, we calibrated and parameterised the models using dynamic quantitative time-course data, and validated them against additional independent datasets. We hypothesized that the most faithful model would be able to fit all the datasets used for training and validation. With this model, we performed specific predictive simulations with an emphasis on the regulation of mTORC1/2 formation and activity, and characterized the governing factors through *in silico* network perturbation analysis. Model predictions were experimentally validated using *in vitro* analysis, and integration and interrogation of public cellular and patient data.

Our predictive modelling and experimental validation revealed a hitherto unknown biphasic dependence of mTORC1 activity on the key mTORC2 component Sin1, uncovering an emergent functional linkage between the two mTOR complexes. This non-linear dependence seems to be a robust feature among a broad array of cell types, as model simulations predict its existence despite the presence of heterogeneity in protein expression levels among various cell types. The Sin1-mTORC1 biphasic response may help explain different context-specific biological observations in cells with low or high levels of Sin1. Furthermore, our results demonstrate that mLST8 is required for the assembly and activity of both mTOR complexes, and suggest mLST8 is a viable therapeutic target in cancer, notably breast cancer. Finally, as the specificity of cellular responses to perturbation is encoded by the spatial and temporal dynamics of signalling networks and coordinated by all network components, this study highlights the importance to embrace systems-based approaches where computational models serve as vital tools to explore and provide insights into intricate relationships between cellular perturbation and response. The computational model developed here will provide a useful resource for future studies and modelling efforts.

## RESULTS

### Construction of mechanistic PI3K/mTOR network models incorporating mLST8-mediated switch

To elucidate the functional role of the mLST8 (de)ubiquitination switch in coordinating mTORC1/2 formation and PI3K/mTORC network dynamics, we constructed new mathematical, mechanistic models of this network incorporating the switch regulation. A number of models have previously been developed for the PI3K/mTOR pathway (14). For example, Pezze et al. (2012) presented a model using ordinary differential equations (ODEs) to investigate potential regulators of mTORC2. Based on observations that amino acids can also activate mTORC2 in addition to mTORC1 (15), an integrated modelling-experimental approach was employed to identify novel downstream targets of amino acids within the mTOR pathway (16). Other models focused on insulin resistance and aimed to explain the PI3K/mTOR network dynamics following insulin stimulation in healthy and diabetic cells (17-20). We have also constructed mechanistic models that revealed complex emergent dynamical properties brought about by DEPTOR, an endogenous inhibitor of both mTORC1 and 2 (21). However, none of the published models consider the coordination of mTORC1/2 formation by the mLST8 (de)ubiquitination switch (7). The mathematical models developed in this study are the first to explicitly incorporate this switch, in addition to known regulatory and feedback mechanisms. Below we outline the key experimental observations and model assumptions.

### mLST8-mediated switch regulates mTOR complex formation

While mLST8 was identified as a shared component of mTORC1 and 2 more than a decade ago (12, 22), only recently the (de)ubiquitination of mLST8 was found to strongly dictate its binding to mTORC1 and 2 (7). Indicated in Fig 1A, mLST8 can be ubiquitinated by the E3 ligase TRAF2 (TNF receptor associated factor 2) on its WD7 kinase domain, which is also the binding site for Sin1. Consequently, mLST8 ubiquitination prevents Sin1-mLST8 association and thus decreases formation of the mTORC2 complex. However, as Raptor binds mLST8 via its WD6 domain, mLST8 ubiquitination does not affect mLST8-Raptor binding (7). Moreover, reduced Sin1-mLST8 association caused by mLST8 ubiquitination frees more mLST8, which becomes available for binding to Raptor, thereby enhancing mTORC1 formation (7). The ubiquitination of mLST8 is reversed by the deubiquitinating enzyme OTUD7B (OUT domain-containing protein 7B), which catalyses deubiquitination of mLST8 (7) and by doing so, it enhances mLST8’s binding affinity for Sin1 and so mTORC2 formation (Fig 1A). Importantly, insulin acts as a triggering input for OTUD7B as insulin stimulation promotes activation of OTUD7B (7). Collectively, mLST8 (de)ubiquitination functions as a molecular switch, where deubiquitination promotes mTORC2 formation while simultaneously blocking the formation of mTORC1, and vice versa. However, how the mLST8-induced switch interplays with other regulatory mechanisms within the PI3K/mTOR network to orchestrate emergent network behaviour is not known. These mechanistic details are described in our new models, indicated in the model reaction scheme (Fig 1B).

### Other key signalling events and feedback loops

The model schematic in Fig 1B further includes key signalling and feedback events induced by insulin stimulation. Briefly, insulin binds to the insulin receptor (IR), triggering IR dimerization, autophosphorylation and activation (23). Activated IR recruits and phosphorylates insulin receptor substrate (IRS) that leads to PI3K recruitment and activation (23). Activated PI3K in turn phosphorylates phosphatidylinositol (3,4,5)-bisphosphate PIP2 and generates phosphatidylinositol (3,4,5)-trisphosphate PIP3, which recruits the kinase PDK1 to the plasma membrane, which subsequently phosphorylates Akt at Threonine 308 (pAkt T308). To allow model simplification for better tractability without compromising dynamic accuracy, we lumped the IRS → PI3K → PDK1 → pAkt T308 cascade into a single step, IRS → pAkt S308 (Fig 1B).

As an AGC family kinase, Akt requires dual phosphorylation to become fully activated (24). To this end, mTORC2 serves as a second Akt kinase and phosphorylates it at Serine 473 (pAkt S473). We assume that double phosphorylated Akt (ppAkt) can be achieved independently through either pAkt T308 or pAkt S473 first (Fig 1B) (25). Moreover, as pAkt S473 alone possesses relatively much weaker kinase activity compared to pAkt T308 and ppAkt (25), we assumed that phosphorylation of the downstream substrate TSC2 is primarily catalysed by the latter, which acts to inhibit TSC2 and releases its repression on mTORC1 activity. Activated mTORC1 phosphorylates S6K1, which in turn phosphorylates IRS on an inhibitory site S636, forming a well-established negative feedback within the PI3K/mTOR pathway that downregulates the input signal (26). In addition, there exists another negative feedback from mTORC1 to IRS via Grb10 (27, 28) but since this acts in a functionally similar manner as the S6K1 feedback, we included only the S6K1 feedback for simplicity (Fig 1B). Finally, we have previously demonstrated that phosphorylated Akt at T308 promotes mTORC2 activation through phosphorylation of its subunit Sin1, generating an important positive feedback loop between mTORC2 and Akt (5). This novel positive feedback is also captured by our models.

### Construction of multiple network model variants

In addition to the more established signalling events above, there are gaps in our network structure understanding. First, although insulin stimulation was shown to promote OTUD7B activation, it is unclear if this is mediated at the level of the insulin receptor or downstream. Therefore, we constructed two different model variants to examine alternative scenarios: *(i)* in model 1, OTUD7B is regulated directly by IR and thus is not influenced by the S6K1-IRS1 negative feedback loop; and (*ii*) in model 2, OTUD7B is regulated by IRS and therefore under the control of the negative feedback (Fig 1C, left panels). Second, as previously discussed, while mLST8 is critical for mTORC2 kinase activity, evidence regarding whether it is required for mTORC1’s functional activity are conflicting (9, 10, 12, 13). This led us to construct two additional models in order to investigate the role of mLST8 in mTORC1 regulation. Specifically, in models 1-2, we assumed that mLST8 is not required for mTORC1’s function. In contrast, in models 3-4, mLST8 binds to Raptor and forms mTORC1, thus mLST8 is required for mTORC1 formation and activity (Fig 1C, right panels).

In summary, we constructed four model variants differing in specific details pertaining to the regulation of OTUD7B and the role of mLST8 in mTORC1 regulation, which allow us to examine competing hypotheses. The new models are formulated using ODEs that represent biochemical interactions as a series of ordinary differential equations based on established kinetic laws (29). Solving these equations allows us to evaluate changes in concentration (i.e. states) of network component proteins over time. The rates of protein-protein interactions (e.g. association and dissociation reactions) were described by mass-action kinetics, and the rates of enzyme-catalysed reactions (e.g. (de)phosphorylation and (de)ubiquitination) were given by Michaelis-Menten kinetics. Detailed description of the models, including ODE equations, reactions rates and model parameters are given in Supplementary Table S1-3.

### Model validation indicates OTUD7B is governed by the S6K1-IRS negative feedback loop

To confer specificity and predictive power to our models, we performed model calibration (i.e. parameter estimation) using insulin-stimulated time-course experimental data obtained from mouse embryonic fibroblast (MEF) cells previously published in (15), which was the primary cell model used for the characterization of the mLST8-induced switch. The data were quantified using the software ImageJ (30) (Fig S1). Parameter estimation was carried out using an optimization procedure based on a genetic algorithm implemented in MATLAB (Materials and methods-Mathematical modeling). To overcome possible issues with model unidentifiability, which is a common phenomenon in signalling network models, we repeated the parameter estimation process 500 times for each of the four models to obtain multiple optimal parameter sets that fit the data equally well, and utilise all the best-fitted sets collectively for subsequent *in silico* analysis. This ‘ensemble’ approach helps avoid biases associated with single best-fitted sets, and provides more confidence in simulation results.

Simulation results of the four models using the corresponding optimized parameter sets demonstrate all of the models recapitulate the experimental data well (Fig 1D). Next, to further assess the accuracy of these models, we compared model simulations with experimental data that was not used in the calibration process. To this end, we utilized insulin-stimulated time-course data from *Traf2^-/-^* MEF cells where the E3 ligase TRAF2 is silenced (7) (Fig 2, right panels). In addition, the temporal dynamics of mLST8 following insulin stimulation in wild-type (WT) MEF cells was not included in the calibration process and therefore was also used for model validation.

**Fig 2.**
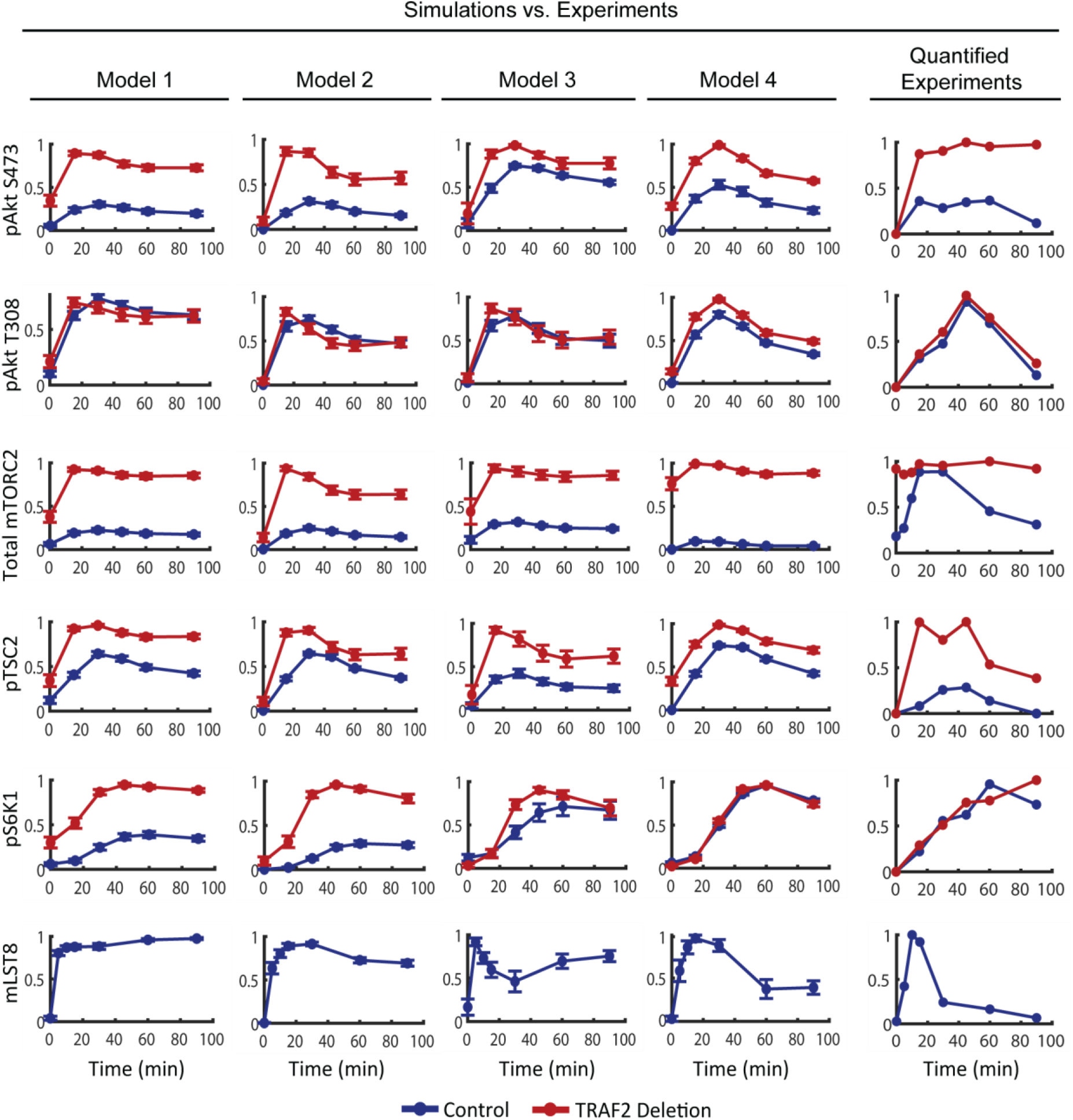
Independent validation of the four models. Simulations of the dynamic response of various network components to insulin stimulation in control (WT MEF cells, blue lines) condition and when TRAF2 is deleted (*traf2*^-/-^ MEF cells, red lines), in comparison to the corresponding experimental data (right panels), taken and quantified from (7). The data are presented as mean ± S.E.M. These validation analysis indicates models 1-3 could not recapitulate all data, and model 4 represents the best-fit model (see text for details).

To mimic TRAF2 silencing, we reduced the concentration of TRAF2 by 90% of its WT level in the models. Comparing model simulations for *Traf2^-/-^* against the WT condition show the models can qualitatively reproduce the temporal dynamics of phosphorylated Akt, TSC2 and mTORC2 (Fig 2). However, model 1, 2 and 3 could not recapitulate the experimentally observed dynamics of phosphorylated S6K1, which was essentially not affected by TRAF2 knockout (Fig 2). In this regard, model 4 is the only model that correctly replicates the data.

In addition, comparing simulated dynamics of mLST8 with the experimental data shows model 1 and 3 failed to capture the transient pattern and low level of mLST8 at late time-points (Fig 2, bottom panels). In contrast, these features are reproduced by model 2 and 4, which differ from model 1/3 in terms of OTUD7B regulation (Fig 1C). In model 2/4, OTUD7B is regulated by the S6K1-IRS negative feedback. Collectively, these *in silico* results indicate OTUD7B activity is governed by the S6K1-IRS negative feedback, and suggest model 4 as the most likely model based on its superior ability to reproduce multiple sets of experimental data.

### mLST8 is required for mTORC1 and mTORC2 activities

To understand why model 4 fits experimental data best, we examined the time-course data more closely. While TRAF2 silencing led to a marked upregulation of both pAkt S473 and pTSC2 (Fig 2 for insulin stimulation, and Fig S2A, B for EGF stimulation), this did not really affect the level of phosphorylated S6K1 (Fig 2 and Fig S2C). This seems counterintuitive since increased pTSC2 will release more mTORC1, resulting in increased mTORC1 activity and phosphorylated S6K1, a direct mTORC1 substrate (see Fig S3A for a visual illustration of this reasoning). This raises the questions as to why pS6K1 was buffered from TRAF2 silencing, and which mechanism underpins such buffering?

We hypothesize that the stable pS6K1 level in response to TRAF2 knockout is due to a compensatory upregulation of mTORC1 activity. Model 4 assumes that mLST8 is required for mTORC1 function. Thus, while there are increased levels of mTORC2 and phosphorylated Akt and TSC2 in *Traf2^-/-^* cells, which together activate mTORC1 more strongly, the abundance of mTORC1 is reduced due to a loss of ubiquitinated mLST8. Overall, this leads to a balance in mTORC1’s total kinase activity potential (defined by the product of mTORC1 abundance and activity potential per mTORC1 molecule, equation (1)), and so no changes in phosphorylated S6K1 (Fig S3B). This balance breaks down in models 1-2, which assume mLST8 is not required for mTORC1 function, resulting in a significant change in pS6K1 when TRAF2 is silenced.

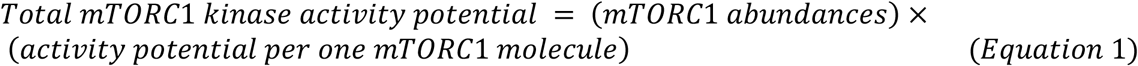

Taken together, through rational construction of a series of mechanistic models and contrasting model simulations with experimental measurements, we arrived at model 4 that could best reconcile multiple sets of experimental data in WT and *Traf2^-/-^* MEF cells. This model will be used for subsequent analyses.

### Modelling predicts biphasic mTORC1 activation dependency on mTORC2 subunit Sin1

Given the key role of mLST8 in coordinating the assembly and activity of mTORC1/2, perturbation of the mLST8-induced switch is likely to disrupt mTORC1/2 signalling but how this occurs is poorly characterised. To address this, we performed *in silico* sensitivity analysis where the concentration of the switch-related proteins (mLST8, TRAF2, OTUD7B, Sin1 and Raptor) were systematically perturbed over a wide range (100-fold up/down) of their nominal values, and the impact on mTORC1 and 2 signalling was quantified using the response of pS6k1 and pAkt S473 at steady state, respectively.

Interestingly, model simulations predict that increasing Sin1 induces a biphasic, dose-dependent response in pS6K1 (Fig 3A). Time-course simulations in Fig 3B confirm that an increase of Sin1 from a low level initially promotes pS6K1 (first phase), but further increase of Sin1 beyond a critical threshold suppresses pS6K1 instead (second phase). Biphasic patterns, although to a lesser extent, are also observed for TRAF2, Raptor and OTUD7B graded perturbations (Fig 3A and Fig S4). The underlying mechanism for Sin1-induced biphasic pattern can be explained based on the fact that mLST8 is competitively sequestered by Raptor and Sin1 for the assembly of mTORC1 and 2, respectively. According to equation (1), the overall ability of mTORC1 to phosphorylate S6K1 is determined by two factors: (i) the abundance of mTORC1 as well as (ii) the kinase activity potential per mTORC1 molecule, the latter is proportionally dependent on the upstream kinases Akt and mTORC2. Thus, the pS6K1 level is dictated by the balance between these factors. During the first phase, an increase in Sin1 would sequester more mLST8 and increase mTORC2 formation, leading to higher mTORC1 activity potential, but at the same time resulting in less mTORC1 formation. As the gain in mTORC1 activity outweighs the loss in abundance, the net effect is an overall enhancement of pS6K1. In contrast, the balance is tipped in an opposite way in the second phase, as a further increase of Sin1 reduces mTORC1 abundance dramatically, which overrides the upregulation in mTORC1 activity, leading to an overall reduction in pS6K1.

**Fig 3.**
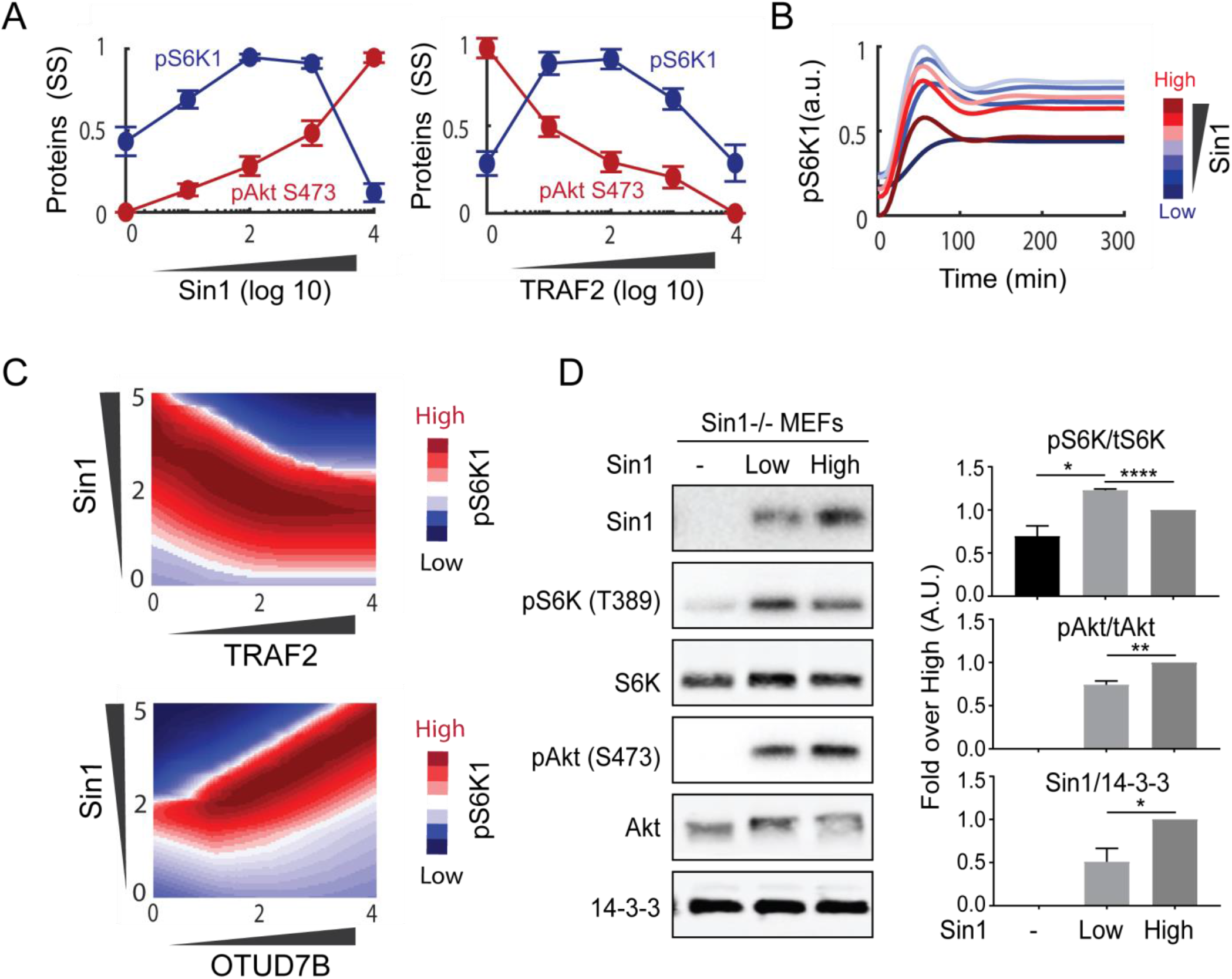
In silico analyses and experimental validation show biphasic dependency of mTORC1 activity on Sin1. (A) Dose-response simulations of the steady-state values of pS6K1 and pAkt473 in response to increasing Sin1 or TRAF2, revealing a biphasic dependence of mTORC1 activity on Sin1. Sin1 or TRAF2 concentrations were varied within a wide range: 100 folds down/up of their nominal values. Each curve related to each parameter set was normalized by its maximal (peak) value and then average and S.E.M. were calculated at each concentration. (B) Temporal simulations of pS6K1 at different concentrations of Sin1. (C) Two-dimensional perturbation analysis showing the simulated steady-state response of pS6K1 to simultaneous change in pairs of proteins, which demonstrates the biphasic pattern is robust over wide ranges of TRAF2 and OTUD7B expression levels. (D) Experimental validation of the Sin1-mTORC1 biphasic dependency. Cells were deprived of serum for 2 h, followed by insulin (100 nM) stimulation for 10 min, and samples were analyzed by Western blot. Graphs show mean and SEM of quantitative analyses of Western blots (*P<0.05, **P<0.01, ****P<0.0001, two-tailed student’s t test, n = 3 biological replicates).

To further investigate the robustness of this biphasic dependency with regard to changes in the expression of multiple proteins, we extended the sensitivity analysis to two dimensions by assessing the level of pS6K in response to simultaneous changes in abundance of pairs of state variables. Fig 3C shows that the Sin1-dependent pS6K1 biphasic response is present over a large range of TRAF2 expression levels. Similarly, the biphasic response persists over a wide range of OTUD7B expression levels. Together, these simulation results indicate that the biphasic mTORC1 activation induced by Sin1 is robust to expression variation in multiple network components.

Unlike pS6K1, model simulation predicts a consistent, monotonic increase of pAkt S473 level in response to overexpression of Sin1 (Fig 3A). Similarly, increasing OTUD7B and TRAF2 concentration monotonically promotes and diminishes pAkt S473, respectively; owing to their opposing effects on deubiquitinated mLST8 and mTORC2 formation (Fig 3A and Fig S4). Interestingly, simulations show that overexpression of mLST8 enhances the level of both pAkt S473 and pS6K1 (Fig S4), suggesting mLST8 promotes activation of both mTOR complexes. The model prediction is consistent with the finding that mLST8 is upregulated in several cancer types (13). mLST8 knock down has beenshown to suppress tumor growth by inhibiting mTOR complex formation and activity (13).

### Experimental validation of Sin1-induced biphasic mTORC1 dependency

In order to experimentally validate the predicted mTORC1 biphasic dependency on Sin1, we generated MEF cells that express increasing levels of Sin1. To this end, we utilized MEFs where Sin1 has been knocked out, and transfected these with the Sin1 construct containing EGFP as a sorting marker. The Sin1 low- and high-expression cells were sorted according to the expression level of EGFP.. To measure the effect of various Sin1 levels on phosphorylated S6K1, cells were stimulated with insulin after serum starvation and pS6K1 was measured using Western Blots (Fig 3D). In cells with no Sin1, there was a low level of pS6K1, resulting primarily from pAkt T308 activity alone since mTORC2 is not functional in these cells, evident by the lack of any pAkt S473 signal (Fig 3D). In cells with low Sin1, there was a significant increase in the level of both pS6K1 and pAkt S473, the latter due to the formation of functional mTORC2. However, in cells with the highest level of Sin1, while the level of pAkt S473 was further increased, the level of pS6K1 was instead significantly reduced in comparison to cells with low Sin1 (Fig 3D). These data clearly confirm model predictions and demonstrate a biphasic pattern in mTORC1 activity in response to graded increases of Sin1 expression. Furthermore, in contrast to the results in previous studies indicating Sin1 knockout and mTORC2 activity have no effects on mTORC1 function (9, 10), our model simulations verified by experimental data here show that mTORC2 regulates both the activity of AKT and mTORC1 in MEF cells.

### Interrogating the Sin1-mTORC1 biphasic dependency in diverse cancer cell lines

By integrating model-based simulation and biological validation, we have identified a previously unknown biphasic dependency between Sin1 and mTORC1 activity in MEF cells. To investigate the existence of this biphasic connection under a wide range of cellular contexts, and how it may be impacted by cell-to-cell variability in protein expression, we adjusted our model by incorporating cell-type specific expression of the model proteins from a diverse array of cell types and performed simulations under these different conditions.

To this end, we first obtained *relative* protein expression recently reported by the Cancer Cell Line Encyclopedia (CCLE) for 375 human cancer cell lines (31), which allowed us to compare protein levels *across* the various cell lines. Of these, 33 cell lines have missing data, i.e. no detectable expression of one of the model proteins, leaving 342 for further analysis. However, tailoring our model to a new cell type ideally requires *absolute* protein levels, i.e. abundance of proteins *within* a proteome. To address this, we utilized a second dataset containing absolute protein abundances obtained by Geiger et al. (32) for 11 common cell lines, using mass spectrometry based label-free proteomics and intensity-based absolute quantification (iBAQ) algorithm (32). Since 7 cell lines were consistent between these two datasets, we could use the iBAQ data of these 7 cell lines and the relative protein information in 342 cell lines to infer the absolute protein levels for the CCLE cohort. A schematic of this inference pipeline is illustrated in Fig 4A. Using MCF7 as an example, we combined absolute protein abundances (i.e. iBAQ data) in MCF7 and the relative expression data (i.e. CCLE data) between MCF7 and 341 remaining cell lines to compute the absolute protein abundances for these cells. We repeated this process for the 6 remaining cell lines with iBAQ data (Fig 4A). As a result, for each of the 342 CCLE cell lines we have 7 *sets* of protein abundances that were inferred using each of the 7 cell lines from (31). The final abundance of the model protein components in each cell line were then derived by taking average of the 7 corresponding values, which were subsequently used to modify the initial conditions in our model in order to tailor it for each of the 342 cancer cell lines.

**Fig 4.**
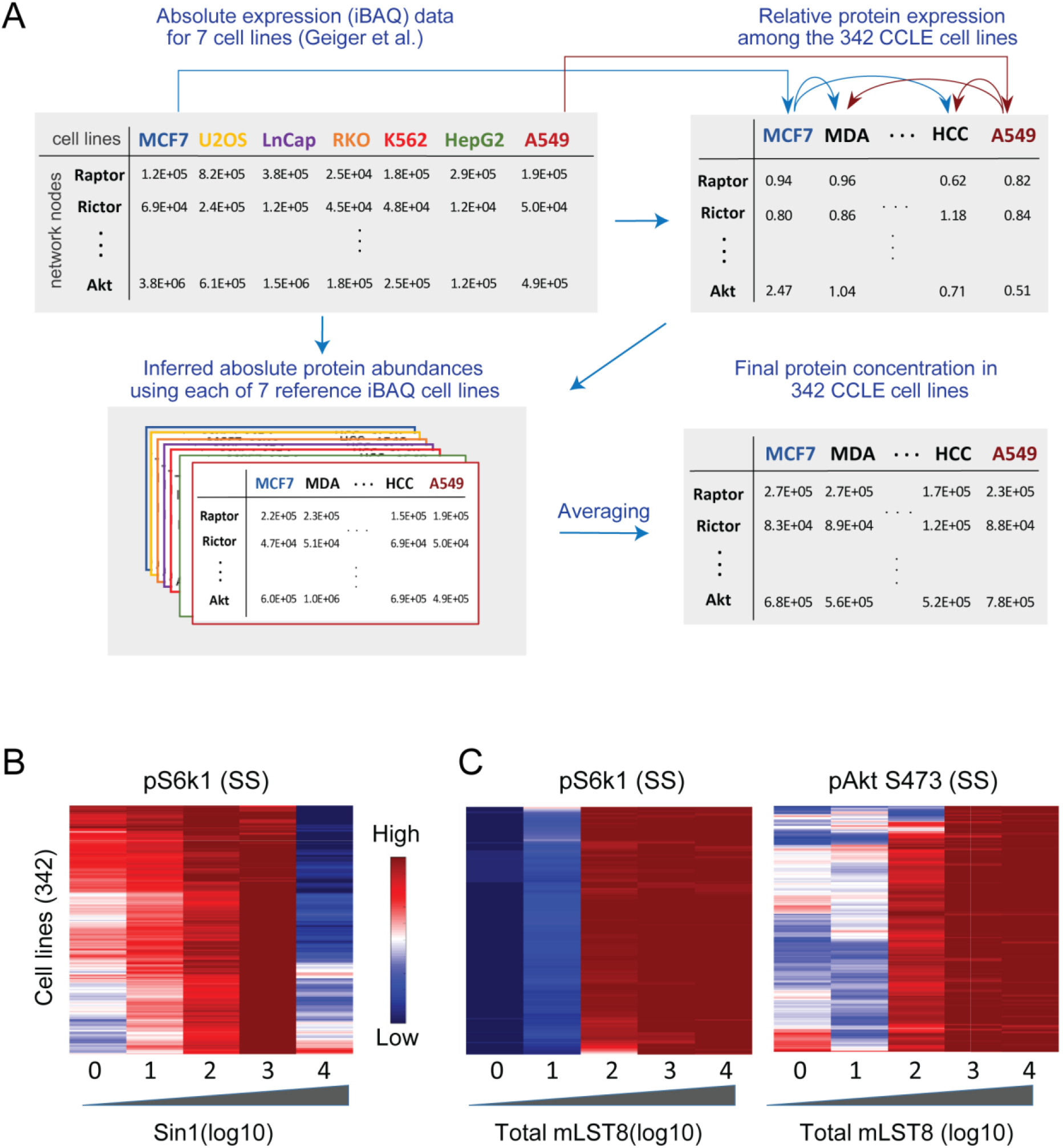
The mTORC1-Sin1 biphasic dependency is robust under diverse cellular contexts. (A) Pipeline for inference of absolute protein levels in 342 CCLE cell lines by combining CCLE relative expression and Geiger iBAQ-based absolute expression data. (B) Heatmap representing dose-response simulation of pS6K1 in response to increasing Sin1 expression, showing the Sin1-mTORC1 dependency is a robust biphasic pattern across multiple cell lines. For each line, Sin1 concentration is perturbed from 0.01 to 100 fold of its initial value and pS6K1 is measured at the steady state (SS). (C) Heatmaps representing dose-response simulation of pS6K1 (left) and pAkt473 (right) in response to increasing mLST8 expression in 342 cancer cell lines.

Having customized our model for different cell lines, we asked if the Sin1-mTORC1 biphasic dependency may still exist under these varying cellular contexts. Fig 4B displays the model simulation results for pS6K1 in response to increasing Sin1 expression, showing the biphasic dependency is still present in almost all of the cell lines. This suggests the biphasic dependency is robust to intercellular variations, albeit the precise shape of this biphasic curve slightly differs between different lines, where it peaks at a lower level of Sin1 in some compared to others (Fig 4B). In addition, we analyzed the effect of mLST8 perturbation in various network conditions (Fig 4C). In contrast to Sin1, increasing mLST8 consistently enhanced the level of phosphorylatedS6K1 and Akt in the majority of the tested cell lines. These results highlight the role of mLST8 overexpression in tumor formation and progression in part through promoting mTORC1 and 2 activities, which has been observed in human colon carcinoma and prostate cancer (13).

### Molecular factors governing the Sin1-mTORC1 biphasic dependency

A strength of a computational modelling approach is that factors controlling emergent complex network behaviors can be identified through systematic *in silico* perturbation/sensitivity analysis, which would otherwise be challenging experimentally. Here, we seek to decipher the molecular players that govern the observed Sin1-mTORC1 biphasic dependency. To evaluate a biphasic response quantitatively, we introduced a general ‘biphasic index’ (BI) that measures the *biphasicness* of a response curve, defined as in Fig 5A. Since the response curve is normalized to its peak (maximal) value, BI ranges between −1 and 1. BI = −1 or 1 indicate a strictly monotonic increasing and decreasing response curve, respectively; while −1< BI <1 indicates biphasic pattern which becomes more pronounced as BI is closer to 0 (Fig 5A). Next, we systematically perturbed the model kinetic parameters representing the strength of network interactions one by one within wide ranges (1000 folds up/down of nominal values), and assessed the impact of these perturbations on the BI of the Sin1-mTORC1 response curve.

**Fig 5.**
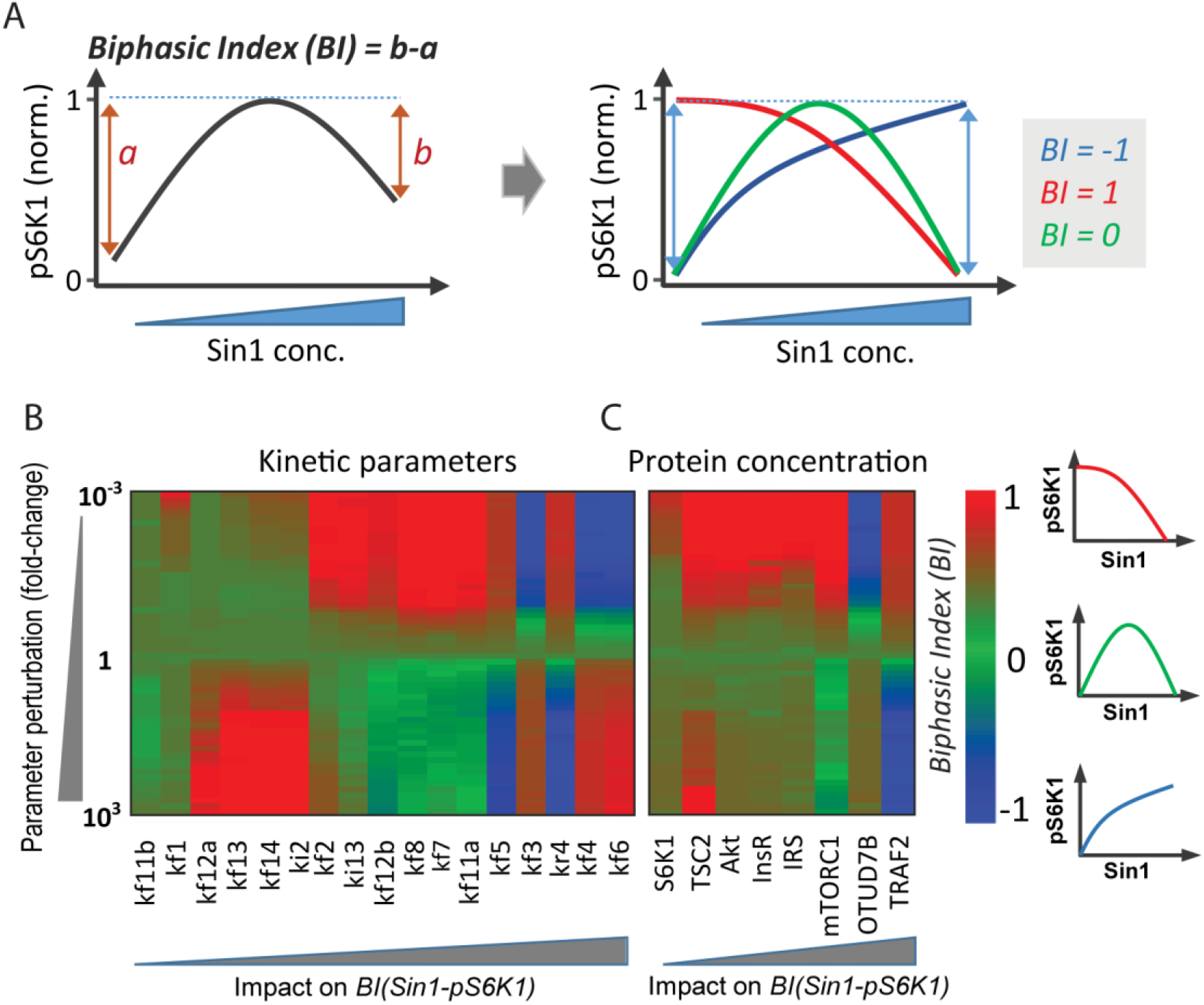
Deciphering the molecular players that govern the Sin1-mTORC1 biphasic dependency. (A) Derivation of a quantitative Biphasic Index (BI) measuring the *biphasicness* of pS6K1 response curve to Sin1 perturbation. The response curve is normalized to its peak (max) value. BI ranges between −1 and 1: BI = −1 or 1 indicate a strictly monotonic increasing (blue) and decreasing response curve (red), respectively; while −1< BI <1 indicates a biphasic pattern that is more pronounced as BI is closer to 0 (green). (B-C) Impact of perturbation of model kinetic parameters (B) or protein concentrations (C) on the BI of the Sin1-mTORC1 response curve. The kinetic parameters were perturbed one by one within wide ranges (1000 folds up/down of nominal values), and the impact on the BI were computed. The parameters/proteins are ranked as having minimum (left) to maximum (right) effect on variation of BI (Sin1-mTORC1).

The results, displayed in Fig 5B, indicate that the parameters related to the mLST8 switch: (de)ubiquitination of mLST8 (*k_f4_* and *k_r4_*), mLST8-Sin1/Raptor binding (*k_f6_*/*k_f5_*), and OTUD7B activation (*k_f3_*) have the strongest impact on the shape of the Sin1-pS6K response curve. Indeed, lowering *k_f3-6_* converts the Sin1-pS6K curve from biphasic to monotonic increasing, while raising them shifts the curve to a monotonic decreasing pattern instead (the opposite is true for *k_r4_*). Consequently, biphasic response exists over relatively restricted ranges of these parameters (green regions, Fig 5B). In contrast, the biphasic response persists over much wider ranges of the lower-ranked parameters, and their perturbations largely shifts the curve an increasing pattern only (green to red, Fig 5B), suggesting the biphasic response is less sensitive to changes in the switch non-related parameters. Of note, the lowest-ranked parameters (*k_f11b_, k_f1_*) do not significantly affect the biphasic pattern. Interestingly, the sensitivity analysis further reveals that *k_f12a_* and *k_f12b_* impact the BI in opposite ways, indicating the rate of TSC2 phosphorylation by pAkt T308 (*k_f12a_*) or the fully activated pAkt S473T308 (*k_f12b_*) have divergent influence on the Sin1-pS6K response. The reason for this is that pAkt S473T308 is regulated by mTORC2 activity and therefore Sin1 concentration, whereas pAkt T308 is independent from Sin1.

In addition, we performed similar sensitivity analysis for the model state variables (i.e. proteins’ concentration). Fig 5C shows that in accordance with the results above, perturbing OTUD7B and TRAF2, the primary regulators of the mLST8 (de)ubiquitination switch, most strongly affect the Sin1-pS6K response BI. Collectively, these results indicate that the mLST8 ubiquitination switch and its constituents critically governs the biphasic relationship between mTORC1 activity and Sin1 concentration.

### mLST8 represents a viable therapeutic target in breast cancer

Our model simulation showed that mLST8 enhances the activity of both mTORC1 and 2 (Fig 4C and S4), suggesting it plays a tumour-promoting role. To examine this further, we interrogated the alteration profiles of mLST8 in patient cancers from The Cancer Genome Atlas (TCGA). In line with our prediction, mLST8 is primarily amplified in the top frequently altered tumour types, including breast cancer, uterine carcinosarcoma and adrenocortical carcinoma (Fig 6A). Further analysis of breast cancer, where mLST8 is most commonly amplified, shows that it is frequently overexpressed (~14%) in patients from both the TCGA and METABRIC cohorts (Fig 6B), two of the largest breast cancer cohorts publicly available. Importantly, our survival analysis demonstrates that high mLST8 expression is associated with worse overall survival in breast cancer patients (Fig 6C). Together, these results indicate mLST8 plays a tumour-enhancing role in breast cancer and its expression may serve as a prognosis indicator, in line with its reported role in other tumour types (12). The findings also points to the exciting possibility of targeting mLST8 as a future therapeutic intervention strategy in breast cancer.

**Fig 6.**
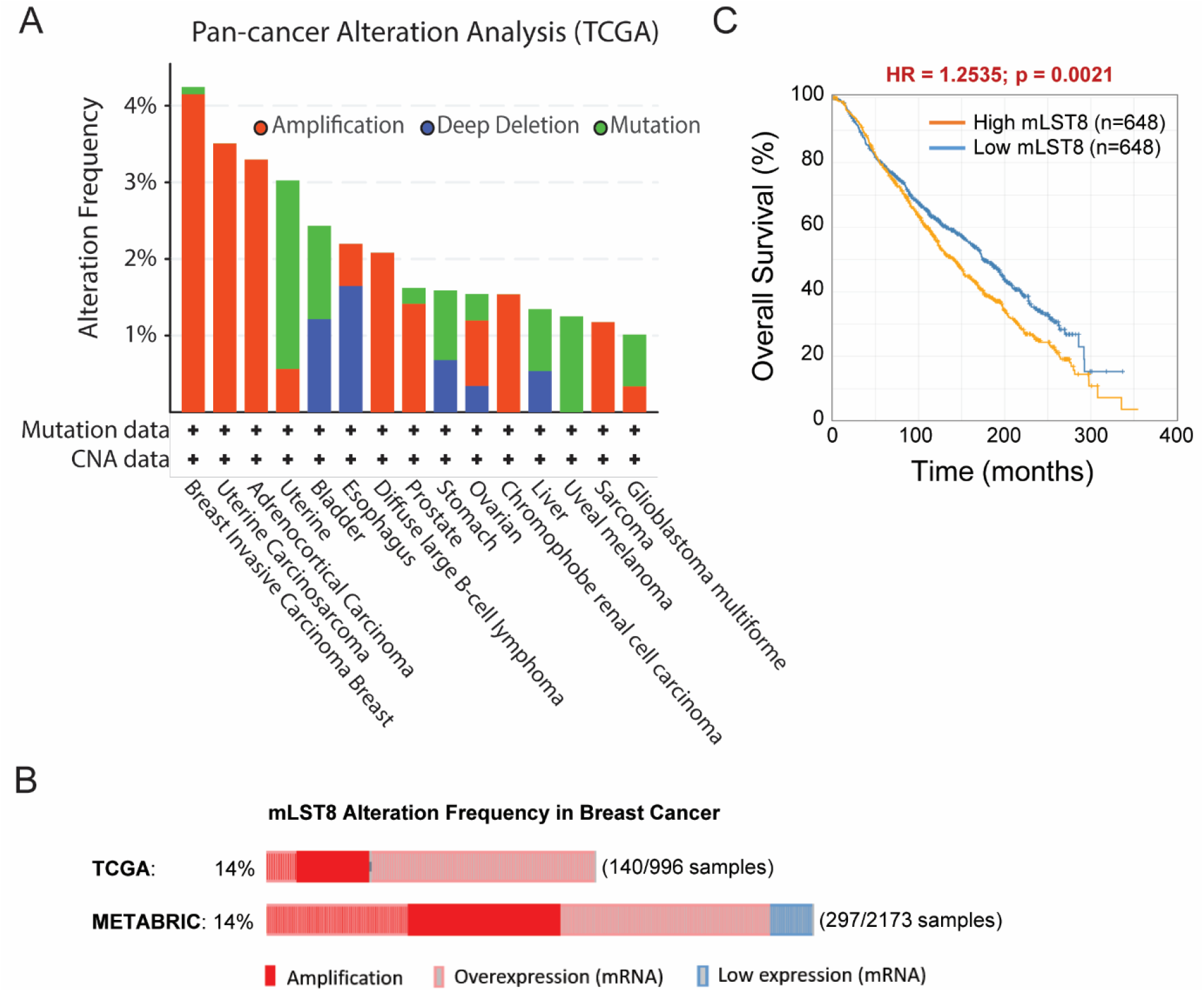
Interrogation of mLST8 alteration and prognostic value in cancer patients. (A) Alteration frequency of the mLST8 gene in cancer patients analysed from TCGA database using cBioPortal (www.cbioportal.org), shown for the most frequently altered tumour types (cut-off > 1%). (B) Frequency of mLST8 alteration in breast cancer patients analysed using two breast cancer cohorts TCGA and METABRIC. Only patients having mLST8 alterations are displayed for brevity. (C) Breast cancer patients with high mLST8 expression are significantly associated with poorer overall survival outcome compared to those with low mLST8 (analysed using the METABRIC cohort data obtained through cBioPortal).

## DISCUSSION

Signalling networks are the key information-processing machineries that underpin the ability of living cells to respond proportionately to extra- (and intra-) cellular cues. More detailed analyses of these networks reveals they are complex interaction systems, often featuring multiple feedback mechanisms, ubiquitous post-translational modifications and intricate pathway crosstalk. In-depth understanding of these networks thus requires quantitative approaches that both complement and extend biological experimentation. We integrate computational network modelling and experimental studies in a systems-based framework to characterize emergent properties of the PI3K/mTOR signalling network, a central regulator of cellular growth, aging and metabolism.

Dynamic modelling has been employed extensively in the study of cell signalling networks, uncovering valuable insights into cellular behavior (33). Here, we present new dynamic models of the PI3K/mTOR network that explicitly accounts for the critical coordination of mTORC1 and 2 formation and function by their shared subunit mLST8, through its (de)ubiquitination reactions (7). To our knowledge, these are the first models to do so. We initially postulated four different network structures and corresponding model variants (models 1-4) that reflect competing hypotheses on OTUD7B regulation and whether mLST8 is essential for mTORC1 functional activity. The latter remains a contentious point among several published studies (9-13). By validating model simulations against multiple existing datasets, we were able to discriminate among the alternative network structures and arrived at model 4 as the most likely model as it provides the strongest fit of these data. Moreover, this model was further validated using biology experiments. The results of these analyses indicated that mLST8 is required for mTORC1 assembly and activity, and the deubiquitinase OTUD7B is stimulated through IRS rather than IR directly.

Using model 4 for subsequent analysis, model simulations predicted a hitherto unknown biphasic dependency between mTORC1 activity and Sin1, a central subunit of mTORC2. Importantly, we confirmed this prediction experimentally by gradually overexpressing Sin1 in Sin1-/-MEF cells, and measured phosphorylated S6K1 and Akt (pS473) as indicators of mTORC1 and 2 activity, respectively. Indeed, increasing Sin1 concentration from a low level significantly enhanced mTORC1 activity, However, beyond a critical threshold, further overexpression of Sin1 inhibited mTORC1 activity (Fig 3A, D). This is in contrast with a monotonic increase in mTORC2 activation in response to Sin1 overexpression observed in both model simulations and experimental data. To identify the molecular factors that control the Sin1-mTORC1 biphasic response, we performed model-based sensitivity analyses assessing possible effects of changes in model kinetic parameters and state variables on the *biphasicness* of the response curve, quantified by a newly derived metric. Interestingly, we found that the top-ranked, i.e. most dominant, parameters and state variables are primarily involved in the regulation of the mLST8 (de)ubiquitination switch, suggesting this molecular mechanism plays an important role in modulating the biphasic level of the Sin1-mTORC1 curve. In line with this finding, the mechanistic explanation underlying the biphasic response stems from the switch-mediated protein competitions and the balance of such competitions that play out within the network. Increasing Sin1 sequesters mLST8 from Raptor to form more mTORC2 and less mTORC1 at the same time; however higher levels of mTORC2 promotes stronger Akt activation, which converts more mTORC1 molecules from an inactive to an active state. In the first phase, the gain in mTORC1 activity dominates the loss in abundance, and the net effect is an enhancement of pS6K1, while the balance is flipped in the second phase, leading to an overall reduction in pS6K1.

While our integrative analyses demonstrated the biphasic Sin1-mTORC1 dependency using MEF cells as the main experimental model, we wondered whether such response is robust to cell-to-cell variability and if it is conserved under different cellular contexts. Given the relevance of the PI3K/mTORC signalling network in cancer and availability of large-scale protein expression data in cancer cell lines, we integrated relative and absolute protein abundances from complementary proteomics profiling studies and developed a pipeline to infer absolute protein abundances for 342 cancer cell lines within the CCLE consortium. Using these data, we adjusted our model for each individual cell line by changing the total expression of model proteins (i.e. initial conditions) accordingly, and simulated the Sin1-mTORC1 response for each line. Simulation results predict that while the precise shape of the response curve vary between the cell lines, the biphasic pattern is still present in most of the considered lines, suggesting the biphasic dependency is quite robust to the cell-to-cell heterogeneity in protein expression and may be a common feature among many cell lines.

This biphasic, rather than monotonic, pattern may underlie the diverse and inconsistent response of phosphorylated S6K to mTORC2 blockade reported by different experimental studies. For example, blocking mTORC2 reduced pS6K in MCF7 and ZR-75-1 breast cancer cells (34) as well as in non-transformed cells, including 3T3-L1 (25, 35) and HEK-293 cells (3); whereas it enhanced pS6K in lung cancer A549 and prostate cancer PC3 cells (35); and induced no significant changes in pS6K in other tumour cells, including HT29 (36) and U251 (37). Moreover, inconsistency has been observed even in the same cell line by different studies, e.g. as in the case of Hela (3, 15) and MEF cells (9, 10). One reason for these discrepancies may simply be due to the very non-linear nature of biphasic input-output curves, where depending on the starting value of the input (dose) and efficiency of knockdown/inhibition, this could either inhibit, promote or not affect the output (response) (see Fig S6).

In addition to revealing a novel functional connection between mTORC2 and mTORC1, our model simulations further showed that mLST8 promotes the activity of both complexes (Fig 4C and S4). This result is consistent with the alteration profiles found in cancer patients analysed from TCGA, which show mLST8 is primarily amplified and overexpressed in the most frequently altered tumour types, notably breast cancer (Fig 6A, B). Model simulations further suggest that mLST8 may stimulate tumour development and/or progression through promoted activation of both mTOR complexes, a notion in line with published data in colon and prostate cancer (13) and our result that high mLST8 is associated with worse overall survival in breast cancer patients (Fig 6C). Together, these findings point to the possibility of targeting mLST8 as an anti-cancer therapeutic strategy, which is indeed being currently investigated (38).

In conclusion, we have integrated mechanistic modelling and experimental analysis to elucidate novel emergent behavior of the PI3K/mTOR signalling network. In contrast to a commonly held view that mTORC2 lies upstream (as often depicted in biological cartoons) and is a positive regulator of mTORC1, we found that their inter-relation is nonlinear and highly dose-dependent. Our results highlight the need to embrace network-level view and apply integrative computational-experimental approaches to study complex signalling and regulatory networks. Our new dynamic model of the PI3K/mTOR network, to our knowledge, is the first that incorporates the mLST8 ubiquitination switch and its critical coordination of the assembly and activity of different mTOR complexes. As such, it provides a novel quantitative framework of the network that is expected to serve as a useful resource for future studies and modelling efforts.

## MATERIAL & METHODS

### Cell culture and viral transduction

PlatE cells were cultured in DMEM with 10% FBS, 2 mM L-GlutaMAX, 1 μg/ml puromycin and 10 μg/ml blastcidine. Sin1-/- MEFs were cultured in DMEM with 10% FBS, 2 mM L-GlutaMAX, non-essential amino acids, and 1 mM sodium pyrophosphate.

For Sin1 retroviral production, PlatE cells (32) grown in 10 cm dish were transiently transfected using lipofectamine 2000 (Life Technologies) according to manufacturer’s instructions with retroviral vectors pMIG or pMIG-Sin1. Medium was replaced the next day with 6 ml medium per dish. Virus-containing medium was collected after 2 days, followed by filtration using a 0.45 micron filter and used immediately for infection or stored at −80°C.

To generate Sin1 re-expression MEFs, Sin1-/- MEFs were seeded into retroviral-containing media and transduced overnight with 4 μg/ml polybrene, followed by fresh media change the next morning.

The Sin1 low- and high-expression cells were sorted by FACS (FACSAria II) according to the expression level of EGFP.

### Western Blotting

Cells were rinsed twice with ice-cold PBS, solubilized in 2% SDS in PBS, sonicated, and spun at 15,000 3 g for 15 min. Protein content was determined by bicinchoninic acid (BCA) assay. Proteins were separated by SDS-PAGE and transferred to PVDF membranes. The membranes were incubated in a blocking buffer containing 5% skim milk in Tris-buffered saline (TBS) and immunoblotted with the relevant antibodies overnight at 4°C in the blocking buffer containing 5% BSA-0.1% Tween in TBS buffer. After incubation, the membranes were washed and incubated with HRP-labeled secondary antibodies for 1 h and then detected by SuperSignal West Pico Chemiluminescent Substrate. In some cases, IR dye 800-conjugated secondary antibodies were used and then scanned at the 800-nm channels using an Odyssey IR imager. Immunoblots were quantified by Image studio software and statistical significance was assessed using Student’s t test. The following antibodies were used: Anti-Sin1 (Millipore, 07-2276); anti-S6K (CST, 2708); anti-S6K-T389 (CST, 9234); anti-Akt (CST, 4051); anti-Akt-S473 (CST, 4058); anti-14-3-3 (Santa Cruz, sc-629).

### Mathematical Modelling

We constructed four mechanistic models to interrogate different possible network structures of the PI3K/Akt signalling pathway. With these models we examined whether: (*i*) The OTUD7B activity is regulated by the S6K1 negative feedback in the pathway, and (*ii*) mLST8 is required for mTORC1 kinase activity. With these four models we assessed the 4 different possible combinations.

The models are constructed using ordinary differential equations (ODEs). The ODEs and the best-fitted parameter sets for each model are given in Tables S1-S3. The model formulation and calibration processes were implemented in MATLAB® (The MathWorks. Inc. 2019a).The IQM toolbox (http://www.intiquan.com/intiquan-tools/) was used to compile the IQM file for a MEX file which makes the simulation much faster.

Model training is the process of estimation of the model’s parameters. As a result of model calibration a ‘best-fitted’ model will be produced that best recapitulates biological data used for model training. To calibrate the model parameters objective function *J* was used that quantifies the difference between the model simulation results and corresponding experimental measurements:

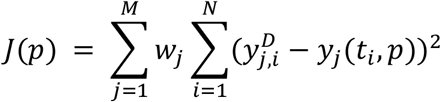

The model parameters value were estimated to minimize the objective function value. Here, *M* is the number of experimental data sets for fitting and *N* denotes the number of time points in a experimental data set. *y_j_*(*t_i_, p*) indicates the model simulations of the component *j* at the time point *t_i_* while parameter set *p* is used for the simulation. Finally, 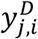 is the experimental data of component *j* at time point *t_i_* and *W_j_* is the weight of the component *j*.

Optimizing the objective function is done by using Genetic Algorithm (GA) (Man et al., 1996; Reali et al., 2017; Shin et al., 2014). For this, Global Optimization Toolbox and the *ga* function in MATLAB were used. To obtain the best fitted parameter sets, GA runs are done with population size of 200 and the generation number set to 800.

### Sensitivity Analysis

To investigate the molecular factors that govern the biphasic pattern between Sin1 concentration and mTORC1 activity we used sensitivity analysis. First, we defined a biphasic index (BI) that calculates the *biphasicness* of the pS6K1-Sin1 concentration response curve. For this, Sin1 concentration is perturbed from 0.01 to 100 of its initial value and pS6K1 is measured to obtain the pS6K1-Sin1 concentration curve (Fig 5A). In the next step, this curve is normalized to 1 and BI index is calculated as defined in Fig 5A.

Next, value of each model kinetic parameter is perturbed from 0.001 to 1000 of its nominal value and the BI is calculated for each parameter value. Finally, the parameters are ranked based on their impact on variation of BI index after perturbing their value.

### Patient Data Analysis

For survival analysis, mRNA expression and associated overall survival data from 2509 breast cancer patients as part of the METABRIC trial (39) were downloaded from the cBioPortal for Cancer Genomics portal (www.cbioportal.org). Patients were classified into groups having low or high expression of mLST8. Analysis comparing overall survival between these groups was done using a Log-rank test (with p< 0.05 considered significant). The Log-rank test statistics and survival curves were generated using Kaplan-Meier estimate and implemented using the Logrank package (https://www.github.com/dnafinder/logrank) using MATLAB 2019b.

## Acknowledgments

The authors acknowledge the facilities, and the scientific and technical assistance at the Biomedicine Discovery Institute, Monash University. The results shown here are in part based upon data generated by the TCGA Research Network: https://www.cancer.gov/tcga.

## Author Contributions

Conceptualization, M.G. and L.K.N; Methodology, M.G., S.S., G.Y., and L.K.N.; Modelling, M.G., S.S., L.K.N.; Experimental Validation, G.Y., D.E.J.; Formal Analysis, M.G., S.S., G.Y. and L.K.N.; Resources, D.E.J. and L.K.N.; Writing (original draft), M.G. and L.K.N.; Supervision, D.E.J., S.S. and L.K.N.; Project Administration, L.K.N.; Funding Acquisition, L.K.N.

## Data and Materials Availability

The reactions, rate equations, differential equations and parameter sets required to reproduce the models can be found in the Supplementary Tables Plasmids generated in this study will be made available upon request. Any further information and requests for resources should be directed to

## SUPPLEMENTARY FIGURES

**Fig S1.**
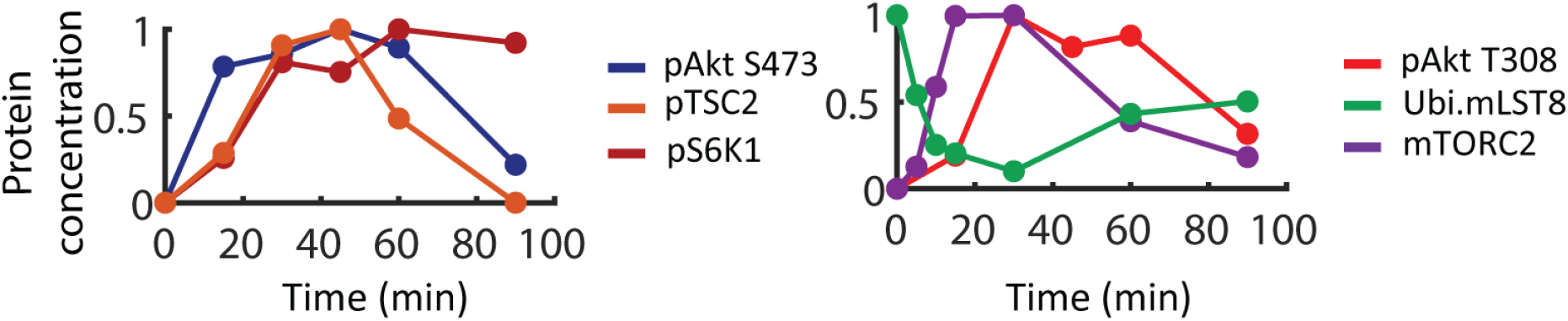
Quantified time-course data of various network components following insulin stimulation (100nM) in WT MEFs, reproduced from ref (7). Phosphorylated (p) and ubiquitinated (Ubi) levels of each protein were normalized to the corresponding total protein levels. Each curve was normalized by their maximal (peak) value.

**Fig S2.**
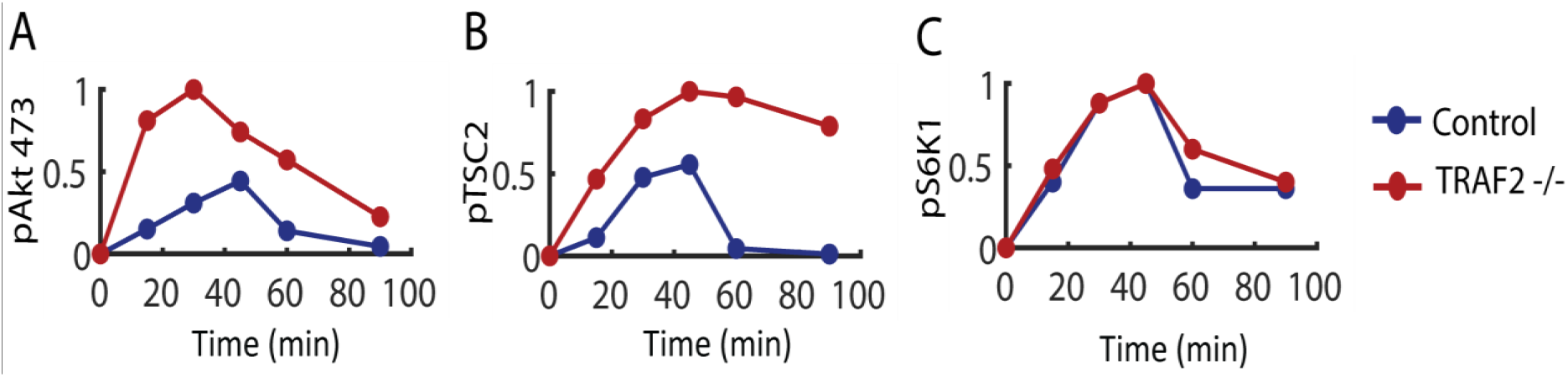
Quantified time-course data of the indicated proteins following EGF stimulation in WT and TRAF2 knocked-out MEF cells, reproduced from ref (7). Phosphorylated (p) levels of each protein were normalized to the corresponding total protein levels. The curves under TRAF2 knockout condition were normalized by their corresponding maximal (peak) values, and the WT curves were scaled accordingly.

**Fig S3.**
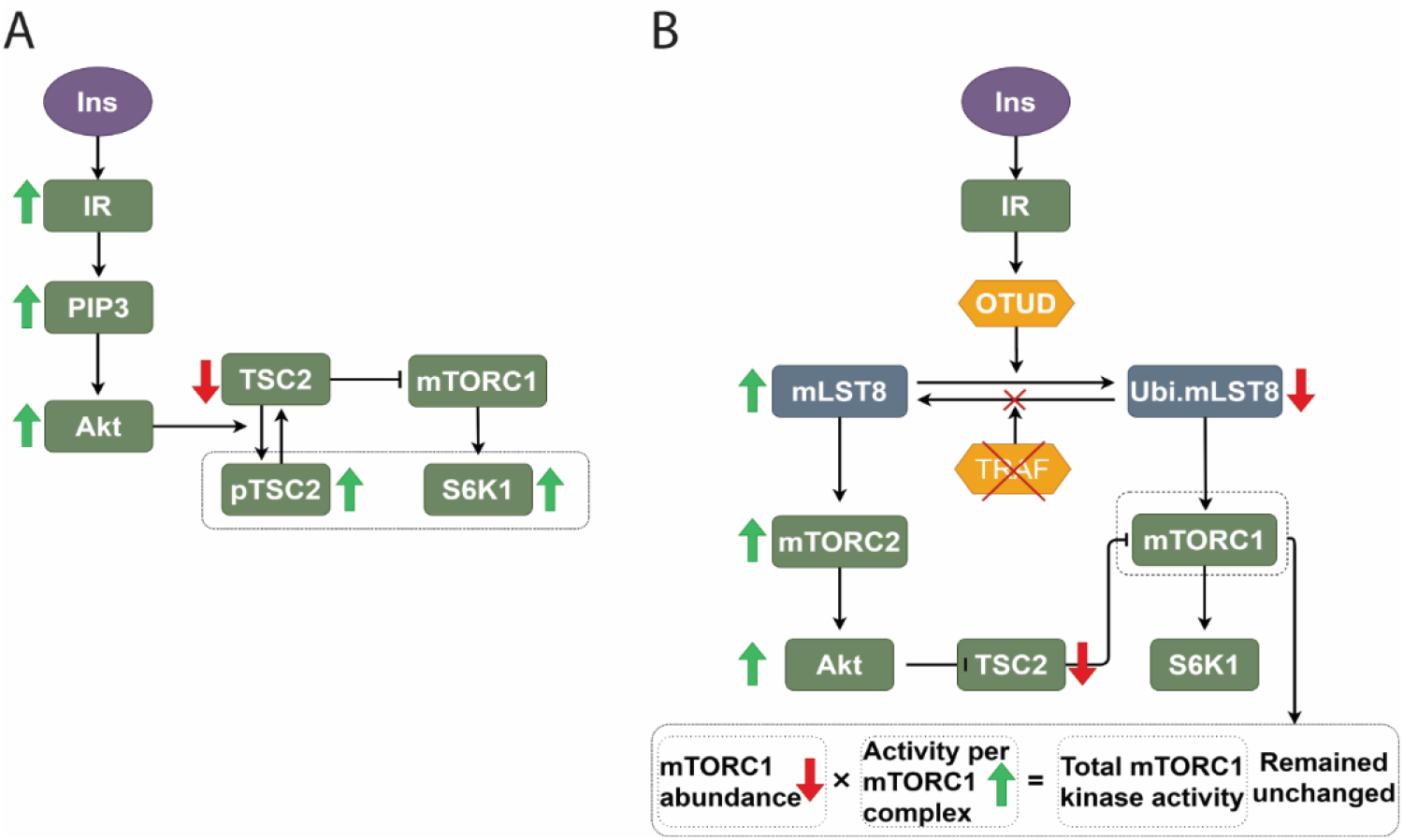
Visual depiction of signal flow within the PI3K/mTOR network. (A) In response to insulin stimulation, phosphorylated Akt increases. This leads to enhanced TSC2 phosphorylation, relief of inhibitory effect on mTORC1, subsequent higher mTORC1 activation and phosphorylation of its key substrate S6K1. (B) Compensatory mechanism following TRAF2 knockout that results in no change in mTORC1 activity dynamics as compared to WT cells: TRAF deletion reduces mTORC1 formation and abundance because of lower ubiquitinated mLST8 but at the same time leads to stronger mTORC1 activity (per molecule) due to higher Akt phosphorylation; and the net effect is no significant change in the phosphorylation of S6K1.

**Fig S4.**
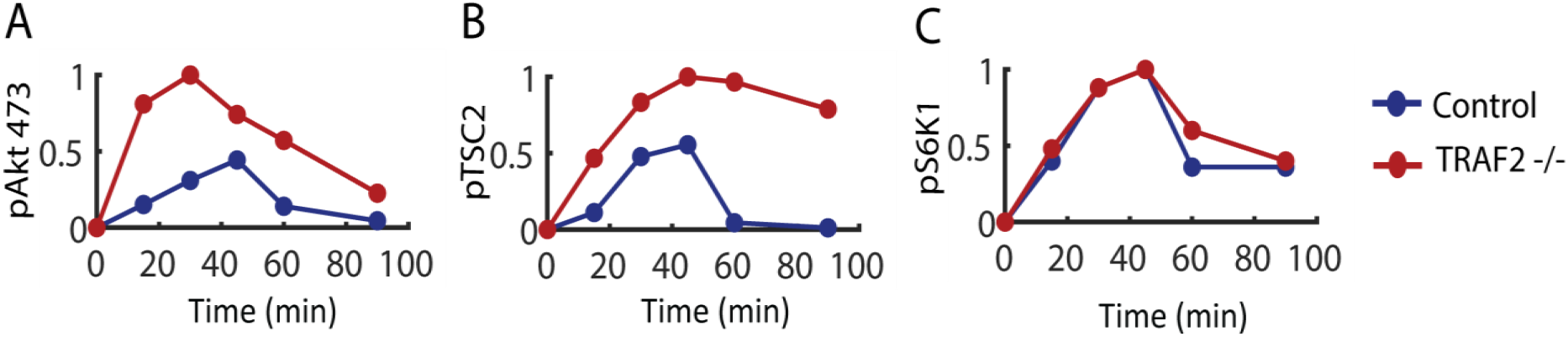
Dose-response simulations of the steady-state values of pS6K1 and pAkt473 in response to increasing Raptor, OTUD7B or mLST8. Total Raptor, OTUD7B and mLST8 concentrations were perturbed within 100 folds up/down of their initial values. The pS6K1 and pAkt473 curves for each parameter set were normalized by their corresponding maximal (peak) values and then average and S.E.M. were calculated at each concentration.

**Fig S5.**
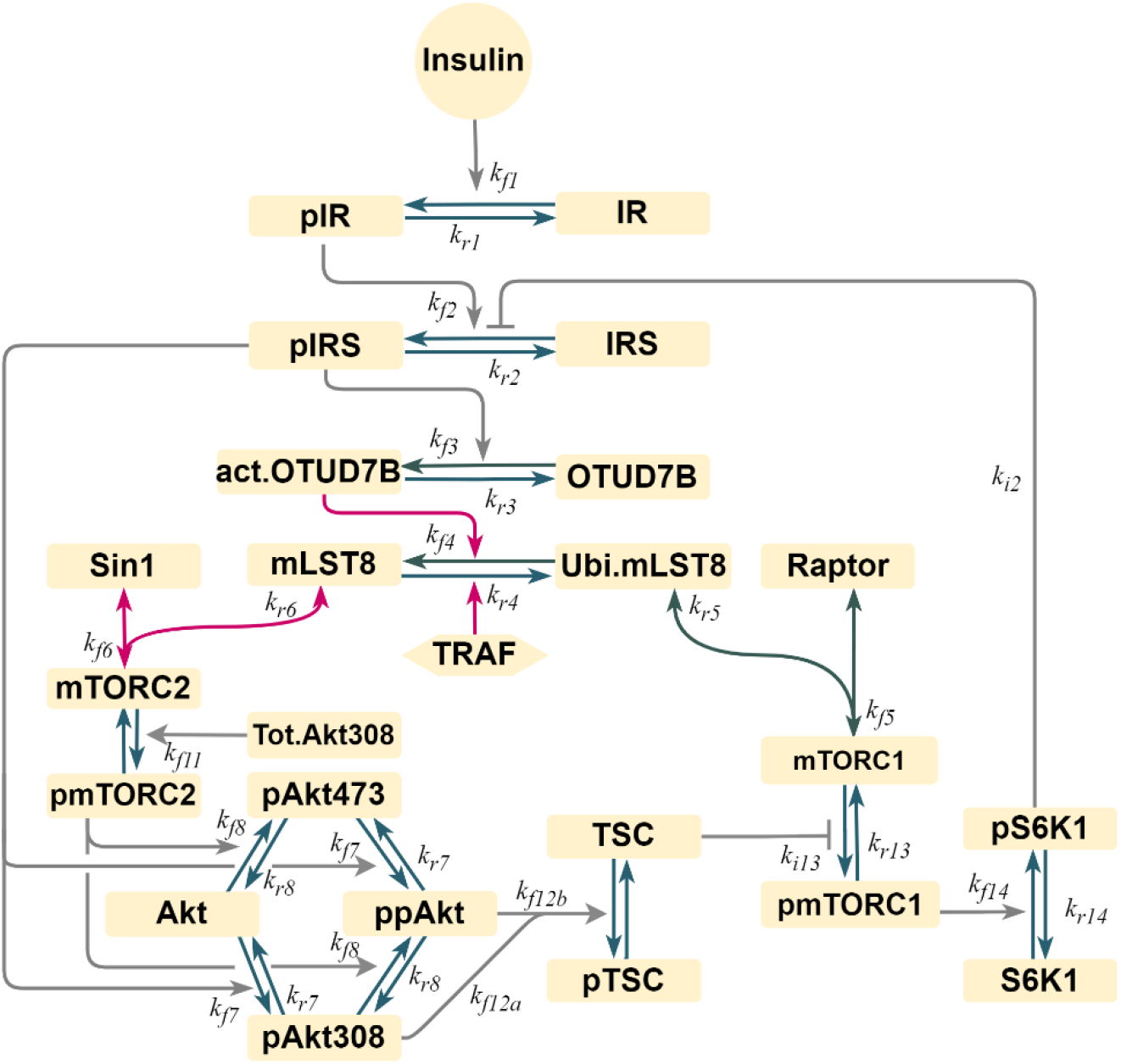
A reaction schematic of model 4 with overlay of the kinetic parameters. Normal arrows indicate positive regulation, bar-headed arrows indicate negative regulation. The red lines indicate the links that exert strongest impacts on the *biphasicness* (i.e. BI) of the Sin1-mTORC1 dependency.

**Fig S6.**
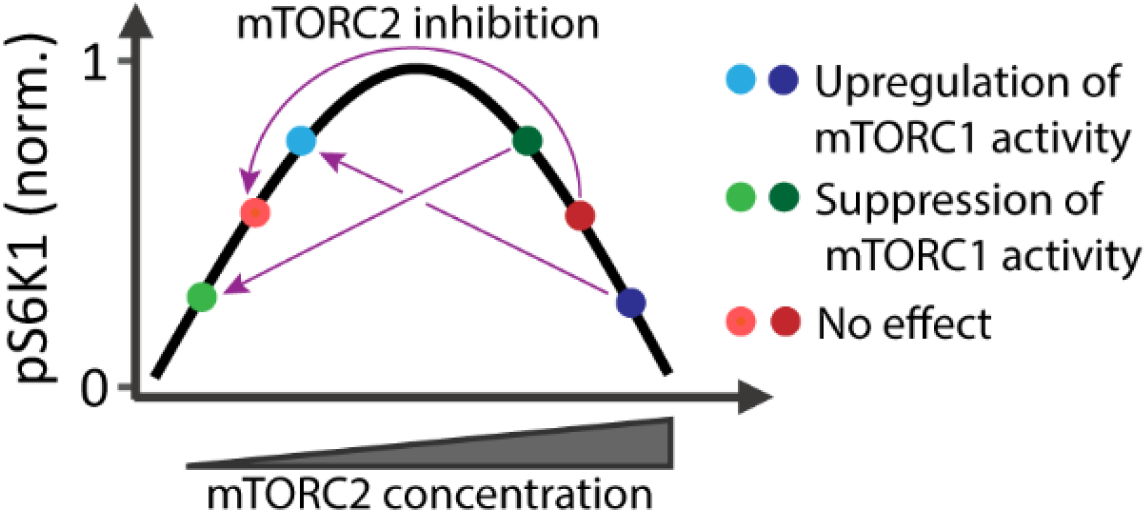
Biphasic response gives rise to highly context-dependent effect of mTORC2 blockade. Depending on the initial state of mTORC2 (e.g. Sin1 level) and the efficiency of blockade, mTORC2 inhibition may result in upregulation, downregulation, or even no change in the phosphorylation level of the key mTORC1 substrate S6K1. This potentially explains the diverse, seemingly conflicting effect of mTORC2 blockade on mTORC1 activity reported in multiple previous studies.

## SUPPLEMENTARY TABLES

**Table S1.** Reactions and rate equations of the PI3K/AKT pathway model

|  | **Reactions** | **Reaction rates** |
| --- | --- | --- |
| R1 | $\text{InsR} \leftrightarrow \text{pInsR}$ | $\text{kf1} * \text{InsR} * (\text{Ins} + \text{InsB0}) / (\text{Km1} + \text{InsR}) - \text{kr1} * \text{pInsR}$ |
| R2 | $\text{IRS} \leftrightarrow \text{pIRS}$ | $\text{kf2} * \text{IRS} * \text{pInsR} / ((\text{Km2} + \text{IRS}) * (1 + \text{ki2} * \text{pS6K1})) - \text{kr2} * \text{pIRS}$ |
| R4 | $\text{Ubi.mLST8} \leftrightarrow \text{mLST8}$ | $\text{kf4} * \text{act.otud} * \text{ubi.mLST8} / (\text{km4} + \text{ubi.mLST8}) - \text{kr4} * \text{mLST8} * \text{TRAF}$ |
| R6 | $\text{mLST8} + \text{Sin1} \leftrightarrow \text{mTORC2}$ | $\text{Kf6} * \text{mLST8} * \text{Sin1} - \text{kr6} * \text{mTORC2}$ |
| R7 | $\text{Akt} \leftrightarrow \text{pAkt308}$ | $\text{kf7} * \text{pIRS} * \text{Akt} / (\text{Km7} + \text{Akt}) - \text{kr7} * \text{pAkt308}$ |
| R8 | $\text{Akt} \leftrightarrow \text{pAkt473}$ | $\text{Kf8} * \text{pmTORC2} * \text{Akt} / (\text{Km8} + \text{Akt}) - \text{kr8} * \text{pAkt473}$ |
| R9 | $\text{pAkt308} \leftrightarrow \text{pAkt308473}$ | $\text{Kf8} * \text{pmTorc2} * \text{pAkt308} / (\text{Km9} + \text{pAkt308}) - \text{kr8} * \text{pAkt308473}$ |
| R10 | $\text{pAkt473} \leftrightarrow \text{pAkt308473}$ | $\text{Kf7} * \text{pIRS} * \text{pAkt473} / (\text{Km10} + \text{pAkt473}) - \text{kr7} * \text{pAkt308473}$ |
| R11 | $\text{mTorc2} \leftrightarrow \text{pmTorc2}$ | $\text{kf11} * \text{mTorc2} * \text{pAkt308} / (\text{Km11} + \text{pAkt308}) - \text{kr11} * \text{pmTorc2}$ |
| R12 | $\text{TSC2} \leftrightarrow \text{pTSC2}$ | $\text{TSC2} * (\text{kf12a} * \text{pAkt308} + \text{kf12b} * \text{pmAkt308473}) / (\text{Km12} + \text{TSC2}) - \text{kr12} * \text{pTSC2}$ |
| R13 | $\text{mTorc1} \leftrightarrow \text{pmTorc1}$ | $\text{kf13} * \text{mTorc1} / (1 + \text{ki13} * \text{TSC2}) - \text{kr13} * \text{pmTorc1}$ |
| R14 | $\text{S6K1} \leftrightarrow \text{pS6K1}$ | $\text{kf14} * \text{mTorc1} * \text{S6K1} / (\text{Km14} + \text{S6K1}) - \text{kr14} * \text{pS6K1}$ |

| **Models** |  | **Reactions** | **Reaction rates** |
| --- | --- | --- | --- |
| Model 1 | R3 | $\text{OTUD7B} \leftrightarrow \text{act.OTUD7B}$ | $\text{kf3} * \text{pInsR} * \text{OTUD7B} / (\text{km3} + \text{OTUD7B}) - \text{kr3} * \text{act.OTUD7B}$ |
| Model 2 | R3 | $\text{OTUD7B} \leftrightarrow \text{act.OTUD7B}$ | $\text{kf3} * \text{pInsR} * \text{OTUD7B} / (\text{km3} + \text{OTUD7B}) - \text{kr3} * \text{act.OTUD7B}$ |
| Model 3 | R3 | $\text{OTUD7B} \leftrightarrow \text{act.OTUD7B}$ | $\text{kf3} * \text{pInsR} * \text{OTUD7B} / (\text{km3} + \text{OTUD7B}) - \text{kr3} * \text{act.OTUD7B}$ |
| | R5 | $\text{Ubi.mLST8} + \text{Raptor} \leftrightarrow \text{mTORC1}$ | $\text{kf5} * \text{Ubi.mLST8} * \text{Raptor} - \text{kr5} * \text{mTORC1}$ |
| Model 4 | R3 | $\text{OTUD7B} \leftrightarrow \text{act.OTUD7B}$ | $\text{kf3} * \text{pInsR} * \text{OTUD7B} / (\text{km3} + \text{OTUD7B}) - \text{kr3} * \text{act.OTUD7B}$ |
| | R5 | $\text{Ubi.mLST8} + \text{Raptor} \leftrightarrow \text{mTORC1}$ | $\text{kf5} * \text{Ubi.mLST8} * \text{Raptor} - \text{kr5} * \text{mTORC1}$ |

**Table S2.** Ordinary differential equations of the PI3K/Akt pathway model. The reaction rates are given in Table S1. The initial conditions are representative values.

| Left hand Sides | Right hand Sides | Initial Conditions (nM) |  |  |  |
| --- | --- | --- | --- | --- | --- |
|  |  | Model 1 | Model 2 | Model 3 | Model 4 |
| $d[\text{InsR}]/dt$ | -R1 | 4.64202 | 55.39126 | 93.71527 | 27.23045 |
| $d[\text{pInsR}]/dt$ | +R1 | 95.35798 | 44.60874 | 6.284733 | 72.76955 |
| $d[\text{IRS}]/dt$ | -R2 | 21.11798 | 99.07924 | 98.72274 | 83.15635 |
| $d[\text{pIRS}]/dt$ | +R2 | 78.88202 | 0.92076 | 1.277256 | 16.84365 |
| $d[\text{OTUD7B}]/dt$ | -R3 | 27.35535 | 99.13060 | 1.069323 | 14.5823 |
| $d[\text{act.OTUD7B}]/dt$ | +R3 | 72.64465 | 0.86940 | 98.93068 | 85.4177 |
| $d[\text{mLST8}]/dt$ | R4 - R6 | 1.29090 | 0.12150 | 1.788563 | 0.017049 |
| $d[\text{Ubi.mLST8}]/dt$ | -R4-R5 | 69.73137 | 99.79011 | 0.05619 | 6.53205 |
| $d[\text{Raptor}]/dt$ | -R5 | — | — | 15.07126 | 16.978 |
| $d[\text{Sin1}]/dt$ | -R6 | 71.02227 | 99.91160 | 86.77349 | 89.5711 |
| $d[\text{mTORC2}]/dt$ | -R11 | 9.06330 | 0.02680 | 13.05844 | 4.215747 |
| $d[\text{pmTORC2}]/dt$ | R11 | 19.91443 | 0.06159 | 0.168063 | 6.213149 |
| $d[\text{mTorc1}]/dt$ | +R5-R13 | 86.74390 | 86.11926 | 84.88032 | 82.782 |
| $d[\text{pmTorc1}]/dt$ | +R13 | 13.25601 | 13.88074 | 0.048424 | 0.240002 |
| $d[\text{Akt}]/dt$ | -R7-R8 | 63.47147 | 99.89068 | 98.82913 | 27.09027 |
| $d[\text{pAkt473}]/dt$ | +R8-R10 | 15.73922 | 0.00153 | 1.90E-05 | 0.828181 |
| $d[\text{pAkt308}]/dt$ | +R7-R9 | 20.75070 | 0.07543 | 1.170799 | 71.97231 |
| $d[\text{pAkt308473}]/dt$ | R9+R10 | 0.03861 | 0.03237 | 5.08E-05 | 0.109242 |
| $d[\text{S6K1}]/dt$ | -R14 | 81.51088 | 79.32881 | 91.89075 | 97.75106 |
| $d[\text{pS6K1}]/dt$ | R14 | 18.48912 | 20.67119 | 8.109249 | 2.248936 |
| $d[\text{TSC2}]/dt$ | -R12 | 35.05656 | 98.55693 | 94.43558 | 21.0847 |
| $d[\text{pTSC2}]/dt$ | R12 | 64.94344 | 1.44307 | 5.564417 | 78.9153 |

**Table S3.** Representative best-fitted parameter sets used for simulations for each of the four models. All the best-fitted parameters are provided in a separated Supplementary Table.

| Parameter | Model 1 | Model 2 | Model 3 | Model 4 | Unit |
| --- | --- | --- | --- | --- | --- |
| Kf1 | 170.2159 | 10000 | 1333.521 | 756.8329 | sec<sup>-1</sup> |
| Km1 | 3.68129 | 0.000931 | 335.7376 | 0.003155 | nM |
| Kr1 | 0.016181 | 1.482518 | 0.028379 | 0.103992 | sec<sup>-1</sup> |
| Kf2 | 90.57326 | 984.0111 | 1025.652 | 31.76874 | sec<sup>-1</sup> |
| Km2 | 4931.738 | 157.3983 | 5794.287 | 0.000369 | nM |
| Ki2 | 2.884032 | 2296.149 | 275.4229 | 42.3643 | nM<sup>-1</sup> |
| Kr2 | 0.075858 | 0.447713 | 0.070469 | 1.425608 | sec<sup>-1</sup> |
| Kf3 | 0.77983 | 4073.803 | 0.952796 | 0.564937 | sec<sup>-1</sup> |
| Km3 | 36.98282 | 2202.927 | 0.010116 | 1.513561 | nM |
| Kr3 | 0.098628 | 255.2701 | 0.535797 | 0.100925 | sec<sup>-1</sup> |
| Kf4 | 47.75293 | 20.18366 | 7.533556 | 4.819478 | sec<sup>-1</sup> |
| Km4 | 54.95409 | 0.389942 | 17.37801 | 0.242661 | nM |
| Kr4 | 138.6756 | 2.218196 | 0.717794 | 232.8091 | sec<sup>-1</sup> |
| Kf5 | – | – | 0.572796 | 3548.134 | nM<sup>-1</sup> sec<sup>-1</sup> |
| Kr5 | – | – | 56.36377 | 4753.352 | sec<sup>-1</sup> |
| Kf6 | 11.27198 | 4.753352 | 1.428894 | 4295.364 | nM<sup>-1</sup> sec<sup>-1</sup> |
| Kr6 | 6683.439 | 3.243396 | 428.5485 | 1555.966 | sec<sup>-1</sup> |
| Kf7 | 58.07644 | 0.001714 | 1169.499 | 5.420009 | sec<sup>-1</sup> |
| Km7 | 215.2782 | 5069.907 | 0.103514 | 8649.679 | nM |
| Kr7 | 6.426877 | 0.037844 | 9015.711 | 0.019634 | sec<sup>-1</sup> |
| Kf8 | 0.474242 | 112.9796 | 1.425608 | 523.6004 | sec<sup>-1</sup> |
| Km8 | 0.299226 | 0.012531 | 0.062806 | 0.002228 | nM |
| Kr8 | 2.697739 | 2.951209 | 0.33266 | 3926.449 | sec<sup>-1</sup> |
| Km9 | 143.2188 | 0.063241 | 1267.652 | 66.06935 | nM |
| Km10 | 0.005129 | 1345.86 | 6280.584 | 475.3352 | nM |
| Kf11a | 0.118577 | 18.96706 | 4.943107 | 46.2381 | sec<sup>-1</sup> |
| Kf11b | 1.883649 | 8912.509 | 0.000935 | 2233.572 | sec<sup>-1</sup> |
| Km11 | 0.003926 | 922.5714 | 5.164164 | 0.103992 | nM |
| Kr11 | 0.271644 | 0.160694 | 0.268534 | 561.048 | sec<sup>-1</sup> |
| Kf12a | 0.003793 | 0.340408 | 2.055891 | 6501.297 | sec<sup>-1</sup> |
| Kf12b | 325.8367 | 4385.307 | 8669.619 | 16.36817 | sec<sup>-1</sup> |
| Km12 | 0.039994 | 255.8586 | 7481.695 | 0.000287 | nM |
| Kr12 | 17.33804 | 0.044875 | 0.007145 | 5929.253 | sec<sup>-1</sup> |
| kf13 | 1054.387 | 0.328095 | 0.366438 | 0.000948 | sec<sup>-1</sup> |
| ki13 | 8016.781 | 16.29296 | 139.9587 | 0.537032 | nM<sup>-1</sup> |
| Kr13 | 0.013428 | 0.025293 | 0.9977 | 0.026546 | sec<sup>-1</sup> |
| kf14 | 0.286418 | 3.614099 | 2904.023 | 35.15604 | sec<sup>-1</sup> |
| km14 | 0.221309 | 0.002009 | 4130.475 | 65.31306 | nM |
| Kr14 | 0.148936 | 1.534617 | 0.058884 | 2.249055 | sec<sup>-1</sup> |
| Ins0 | 1 | 1 | 1 | 1 | nM |

